# Chemoinformatics-guided discovery of food-grade anionic stabilizers for phycocyanin under acidic conditions

**DOI:** 10.64898/2026.05.25.727568

**Authors:** Kevin Chuang, Lorry Luo, Luke Law

## Abstract

Phycocyanin (PC) is the principal natural blue pigment used in functional beverages, but it rapidly loses color and aggregates under acidic conditions (pH ≈ 3). Experimental screening of stabilizers is costly and combinatorially intractable. Here we develop a chemoinformatics framework — descriptor-based QSPR, a chemistry-prior heuristic, and virtual screening — that learns from three rounds of commissioned screening (48 compounds, 6% hit rate) to predict stabilizer efficacy directly from molecular structure. In this genuinely small-data regime (3 positives), a LightGBM classifier built from 10 RDKit descriptors and 11 domain-expert charge/polymer features attained a leave-one-out AUC of 0.73, only marginally above a single-feature charge-density baseline (AUC 0.67); LOO sensitivity was 1/3 at threshold 0.5. A complementary chemistry-prior heuristic encoding anion-type priors substantially outperformed both, reaching AUC 0.95, indicating that explicit chemical knowledge captures information that descriptor-based ML cannot readily recover at this dataset size. SHAP analysis of the QSPR identified effective negative-charge density per unit, log molecular weight, polyphosphate identity, and functional-group density as the dominant features (jointly ≈97% of mean |SHAP|), recovering the electrostatic-complexation mechanism without it being supplied as a prior. Virtual screening of 30 generally recognized as safe (GRAS) food additives nominated the pyrophosphate family — led by sodium pyrophosphate decahydrate and sodium hexametaphosphate (SHMP), both at P ≈ 0.99 — and the heuristic additionally flagged sodium phytate (IP□), which the descriptor model under-ranked at P = 0.045. Experimental validation at pH 3 and 46 °C for 7 days confirmed SHMP 2:1 (78.1 ± 11.3% color retention), TSPP 2:1 (54.1 ± 10.6%) and IP□ 1:1 (52.7 ± 9.0%), while a ternary IP□ + STPP combination reached 83.8 ± 11.9%, surpassing all single-component formulations. ζ-Potential measurements indicated a predominantly electrostatic origin for the protection (Pearson r = −0.82 between ζ and CR□ □ □; n = 24; p = 1 × 10□ □). The framework, dataset and code are released to accelerate stabilizer discovery for other acid-sensitive food colorants and to provide a candid small-data benchmark.

## 1. Introduction

Phycocyanin (PC), a 220-kDa pigment-protein complex extracted from Spirulina, is the only natural-source blue food colorant approved under EFSA, FDA and CFSA for general use in beverages and confectionery [1,2]. Its market is expanding rapidly with the consumer shift away from synthetic dyes such as Brilliant Blue FCF [3]. However, PC is intrinsically labile under the acidic and warm conditions characteristic of soft drinks, sports drinks and dairy beverages (pH 2.5–4, ambient transport at 25–46 °C): below its isoelectric point (pI ≈ 4.6) the protein gains net positive charge, partially unfolds, and aggregates into colorless precipitates within days [4,5]. The phycocyanobilin chromophore further undergoes a conformational transition from an extended to a cyclic form, producing irreversible color loss [6].

Conventional approaches to PC stabilization rely on the addition of anionic or charged polymers — alginate, carrageenan, chitosan, polysaccharides, polyphosphates — selected by laborious screening campaigns [7–9]. The chemical space of food-grade anionic additives is large but irregularly sampled: many GRAS-listed compounds have never been tested, while a small number of polysaccharides have been examined repeatedly. Reported screens are typically dose-response surveys with semi-quantitative read-out (visual □/×), and even within single laboratories, results can vary with batch, storage condition and concentration [10]. The combinatorial cost of expanding such studies to multi-component formulations or to the >10^3^ GRAS additives is prohibitive.

Chemoinformatics has transformed analogous problems in pharmaceutical hit discovery, where quantitative structure-property relationship (QSPR) models trained on hundreds to thousands of compounds enable virtual screening of large candidate libraries [11,12]. Application to food formulation, however, remains rare. The few existing studies focus on flavor- and aroma-compound property prediction or on the physicochemical-property modeling of food-relevant compounds [13,14]; to our knowledge, no published QSPR model targets the stabilization of PC or other acid-sensitive natural colorants. The principal obstacle is dataset size: a typical food laboratory has tested fewer than 100 compounds, whereas conventional QSPR best practice favors datasets of several hundred or more compounds [15].

Recent advances in interpretable machine learning, ensemble methods, and transfer- and low-data learning have re-opened the small-data regime [16,17]. LightGBM with leave-one-out cross-validation and SHAP attribution can build defensible classifiers from tens of examples [18]; chemistry-aware heuristics that encode expert priors over anion class and pKa offer a complementary baseline that does not require descriptor calibration [19]. Combined with virtual screening of curated food-grade libraries, this combination promises an order-of-magnitude reduction in experimental campaign cost.

In this work we (i) consolidate three rounds of commissioned PC stabilizer screening (48 compounds, including 30+ GRAS food additives) into the first structured QSPR-ready dataset for this protein; (ii) train and interpret a LightGBM classifier and compare it against a single-feature electrostatic baseline and a chemistry-prior heuristic; (iii) virtually screen a 30-compound GRAS library to nominate experimentally untested candidates; and (iv) validate three structurally distinct predicted hits experimentally at pH 3 and 46 °C, including a ternary combination that surpasses all single-component formulations. The dataset, model and protocols are publicly released to provide a reusable benchmark for stabilizer discovery across acid-sensitive food colorants.

## 2. Materials and Methods

### 2.1. Materials

Food-grade phycocyanin (E18+, lot PC-E18-260509-B) was obtained from Yunnan Lü-A Bio (Kunming, China). Sodium tripolyphosphate (STPP, CAS 7758-29-4, lot STPP-260428-03; Aladdin), sodium hexametaphosphate (SHMP, 10124-56-8, SHMP-260502-11; Macklin), tetrasodium pyrophosphate (TSPP, 7722-88-5, TSPP-260430-07; Macklin) and sodium phytate (IP□, 14306-25-3, IP6-260421-02; Sigma) were all of food-grade purity. The remaining compounds and their suppliers are listed in Supplementary Table S1. Ultra-pure water (18.2 MΩ·cm) was used for all preparations.

### 2.2 Screening dataset

The training dataset comprises 48 unique compounds aggregated from three rounds of laboratory screening conducted between November 2025 and March 2026. Round 1 examined fucoidan, dextran sulfate sodium (DSS) of varying molecular weight, sodium polyphosphate, sodium metaphosphate, STPP and lignosulfonate at PC ratios from 1/20:1 to 5:1. Round 2 re-screened the same anionic polysaccharides under tightened pH control (pH 3, 7 days) and added the cyclodextrin derivative sugammadex (some Round-1 compounds were therefore tested in both rounds). Round 3 expanded to food-grade sugars (mono-, di-, oligo- and polysaccharides), sugar alcohols, monovalent and polyvalent salts, organic acids, gelatin and acesulfame potassium. Each compound was assigned a binary label: ‘effective’ if it preserved blue color across at least one ratio under the standardized condition (0.375% w/w PC, pH 3, 46 °C, 7 days), and ‘failed’ otherwise. Compounds that produced visible precipitates at the working concentration were labeled ‘failed’. Three compounds met the effective criterion: sodium tripolyphosphate (STPP), sodium polyphosphate and dextran sulfate sodium (MW ≈ 20 kDa).

### 2.3 Molecular descriptors

Each compound was encoded by 21 features. Ten general descriptors were computed with RDKit (MolWt, MolLogP, TPSA, NumHDonors, NumHAcceptors, FormalCharge, RingCount, NumRotatableBonds, FractionCSP3, NumHeavyAtoms). Eleven domain-expert features captured charge chemistry that conventional QSPR misses for ionic food additives: polymer flag, log□ □ (MW), one-hot anion class (phosphate, polyphosphate, sulfate, sulfonate, carboxylate), functional-group density per repeat unit, effective negative charge per unit weighted by anion pKa (polyphosphate 1.0, sulfate 1.0, carboxylate 0.4 at pH 3), and a polyvalent-metal indicator. Polymer compounds with undefined SMILES were assigned default RDKit values based on monomer estimates (Supplementary Methods).

### 2.4 QSPR classifier

A LightGBM gradient-boosted classifier (n_estimators=200, learning_rate=0.04, num_leaves=15, max_depth=4, L2 reg λ=1.0, min_child_samples=2, random_state=20260519) was trained with class weights set to balance the 3:45 positive/negative imbalance (pos_weight = 15). Model performance was evaluated by leave-one-out cross-validation (LOO-CV). For comparison, a baseline univariate logistic regression on effective negative charge density and a chemistry-prior heuristic encoding pKa- and anion-class-aware priors were evaluated under the same LOO scheme. Feature attribution used TreeSHAP [20].

### 2.5 Chemistry-prior heuristic baseline

As a knowledge-based comparator, a deterministic rule set was implemented to assign P(effective) directly from the compound name and anion class without computing descriptors. Polyphosphates (STPP, pyrophosphate, metaphosphate, SHMP family) were assigned P=0.85; non-polyphosphate inorganic phosphates P=0.45; carrageenans P=0.60; other sulfates P=0.50; sulfonates P=0.30; alginate-type carboxylates P=0.35; non-alginate carboxylates P=0.25; phytate P=0.75; cationic polymers P=0.05; otherwise P=0.20. These priors encode published pKa values and reported stabilization activity from the food-hydrocolloid literature. The full rule code is provided as Supplementary File S2.

### 2.6 Virtual screening library

Thirty additional GRAS or food-grade compounds not present in the training set were curated from EFSA E-numbers and the Chinese GB 2760 additive list, prioritizing anionic polymers and phosphates. The feature pipeline of §2.3 was applied identically. Predicted P(effective) was obtained from the final QSPR classifier (retrained on all 48 compounds) and from the chemistry-prior heuristic; the union of top-10 predictions from each method defined the candidate pool for experimental validation.

### 2.7 Validation experiment design

Three predicted hits were selected for experimental validation: sodium hexametaphosphate (SHMP; QSPR rank 2, P=0.99), tetrasodium pyrophosphate (TSPP; QSPR rank 6 within the pyrophosphate cluster, P=0.97), and sodium phytate (IP□; chemistry-prior heuristic P=0.75 — the highest-scoring non-pyrophosphate candidate; QSPR rank 9, P=0.045 — an explicit out-of-distribution test). The Top-3 compounds together with a negative control (PC + HCl) and a positive control (STPP at 2:1) were tested in triplicate at stabilizer:PC mass ratios of 1:1 and 2:1, plus a ternary mixture (IP□ + STPP, 1:1:1). Eight groups × three independent bottles = 24 samples. PC was diluted to 0.375% (w/w), the stabilizer was added at the specified ratio, 1.00 M HCl was titrated dropwise under magnetic stirring (200 rpm) to pH 3.00 ± 0.05, samples were brought to 10.0 mL total volume in 24-mL amber headspace vials, purged with nitrogen for 30 s, sealed, and incubated at 46.0 ± 0.5 °C in the dark for 7 days. Temperature was logged continuously with an Onset HOBO MX1101 logger; the recorded range was 45.5–46.8 °C.

### 2.8 Spectroscopic and ζ-potential measurements

UV-visible spectra (350–750 nm, 1 nm step) were acquired on a Shimadzu UV-2600i spectrophotometer in 1 cm quartz cells at t = 0 and t = 7 d. Color retention was computed per bottle as CR□ □ □ = [A□ □ □ (t=7d) − A□ □ □ (blank)] / [A□ □ □ (t=0) − A□ □ □ (blank)] × 100%, with blanks containing the same stabilizer at the same pH but no PC. ζ-Potential was measured on a Malvern Zetasizer Nano ZS using DTS1070 folded-capillary cells; samples were diluted 1:10 (or 1:20 if PDI exceeded 0.5) in 1 mM NaCl pH 3.00 buffer; each bottle was measured three times within the instrument at 25 °C with 60 s equilibration. Latex standards (target −55 ± 5 mV) were measured every five samples; observed standard values were −53.7 and −56.1 mV. Two ζ measurements with PDI > 0.5 (B14, B21) were excluded and re-acquired under the same protocol per the SOP.

### 2.9 Statistical analysis

All comparisons against the negative control used one-sided Welch’s t-tests (alternative: treatment > G0), reflecting the directional hypothesis that stabilizers should preserve color. Synergy of the ternary combination was assessed by one-sided Welch’s t-test against each single-component arm. Per-bottle correlation between ζ-potential and CR□□□ used Pearson’s r. p-values < 0.05 were considered significant. Holm–Bonferroni adjustment was applied to control the family-wise error rate across the seven vs-G0 contrasts. All statistical computations were performed in SciPy 1.15.3 (Python 3.13).

## 3. Results

### 3.1 The 48-compound iterative dataset

The three rounds of laboratory screening produced a binary-labeled dataset of 48 unique compounds spanning the major food-additive chemical classes — salts, sugars, sugar alcohols, organic acids, polysaccharides, polyphosphates and one protein (Figure 1). The overall hit rate was 6% (3/48); the positive set comprised two polyphosphates (STPP, sodium polyphosphate) and one highly sulfated polysaccharide (dextran sulfate sodium, MW ≈ 20 kDa). Round 3 alone screened 35 generic food additives (sugars, salts, organic acids, sugar alcohols) and produced zero new positives, establishing the chemical scarcity of effective stabilizers in the canonical food-grade space.

**Figure 1.**
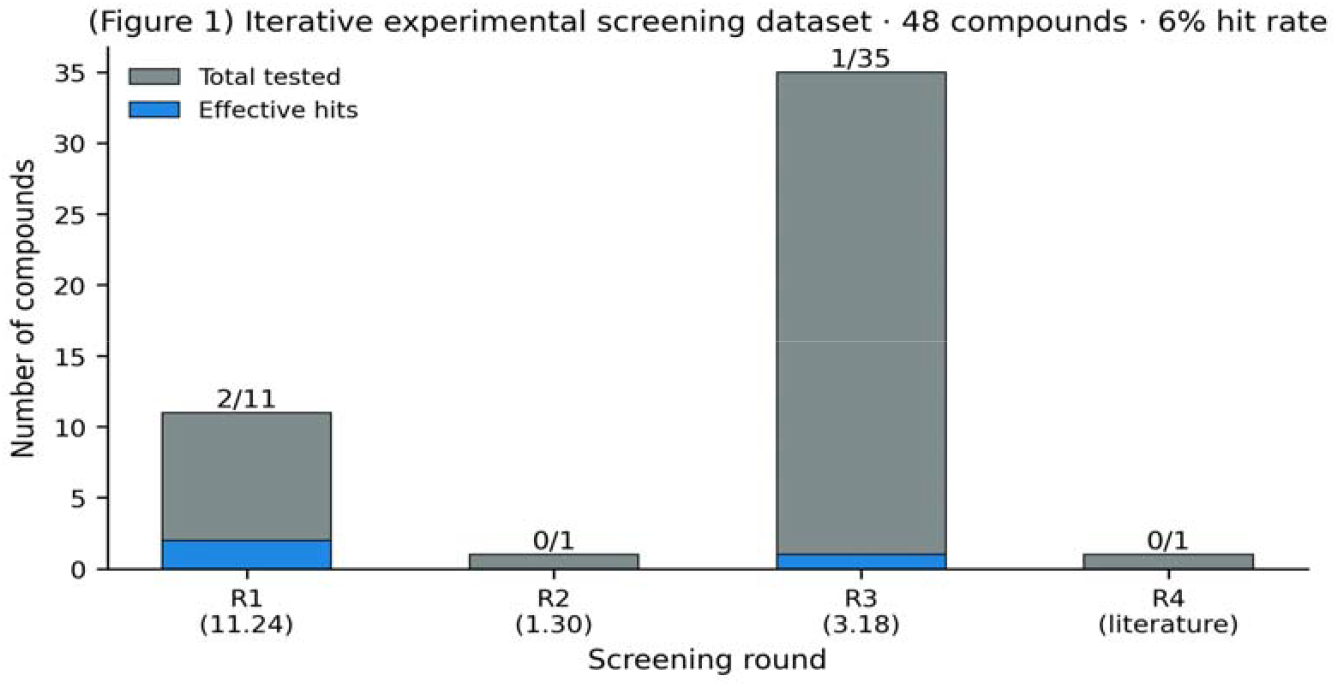
Three-round screening produced a 48-compound dataset with a 6% effective-hit rate. The first two rounds focused on anionic polysaccharides and polyphosphates; the third round broadly sampled food-grade sugars, salts, organic acids and sugar alcohols.

### 3.2 In the small-data regime, descriptor-based QSPR is weak; a chemistry-prior heuristic dominates

The LightGBM classifier reached a leave-one-out AUC of 0.73 (accuracy 0.92), only marginally above a logistic-regression baseline trained on effective negative-charge density alone (AUC 0.67). At a probability threshold of 0.5, LOO sensitivity was 1/3: the model recovered only sodium polyphosphate among the three labeled positives and missed STPP (P = 0.013) and the 20-kDa DSS (P = 0.006), with two false positives (sodium phosphate tribasic, dextran sulfate sodium 15 kDa). With only three positives in the training set, LOO sensitivity is inherently unstable; the AUC reflects ranking quality more reliably than the threshold-0.5 classification. A chemistry-prior heuristic encoding anion class and pKa priors substantially outperformed both descriptor-based models, reaching AUC 0.95 (Figure 3). In this regime, an explicit domain-knowledge baseline therefore provides a more reliable cross-check than descriptor-based ML alone. The descriptor-based QSPR and the rule-based heuristic agreed strongly on the polyphosphate family but diverged on phosphate esters with cyclic scaffolds (Figure 5), foreshadowing the IP□ result reported below.

### 3.3 SHAP attribution recapitulates the electrostatic mechanism

SHAP feature importance (Figure 2) identified, in decreasing order, effective negative-charge density per unit (≈47% of mean |SHAP|), log molecular weight (≈19%), polyphosphate identity (≈19%) and functional-group density (≈12%) as the dominant drivers; these four features jointly account for ≈97% of model attribution. The model thus rediscovered the electrostatic-complexation mechanism — high local negative-charge density at pH 3 produces strong heteroaggregates with the positively-charged PC monomer — without being supplied with this prior, and is qualitatively consistent with published findings on charged-polymer-mediated electrostatic stabilization of PC [7].

**Figure 2.**
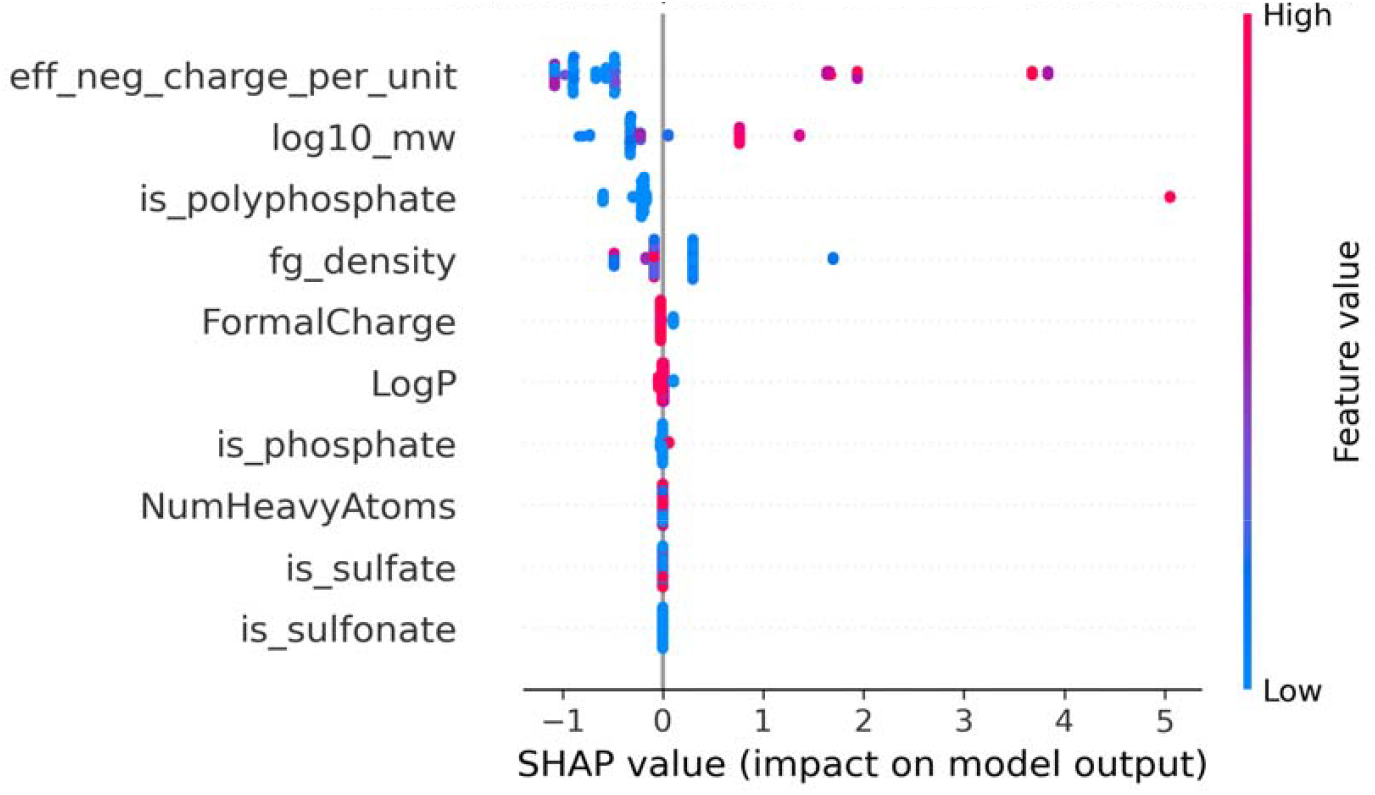
SHAP summary plot. Each point is a compound; horizontal position is the contribution to model output; color encodes the feature value (red = high, blue = low). The pattern recovers the known electrostatic protection mechanism without supplying it as a prior.

**Figure 3.**
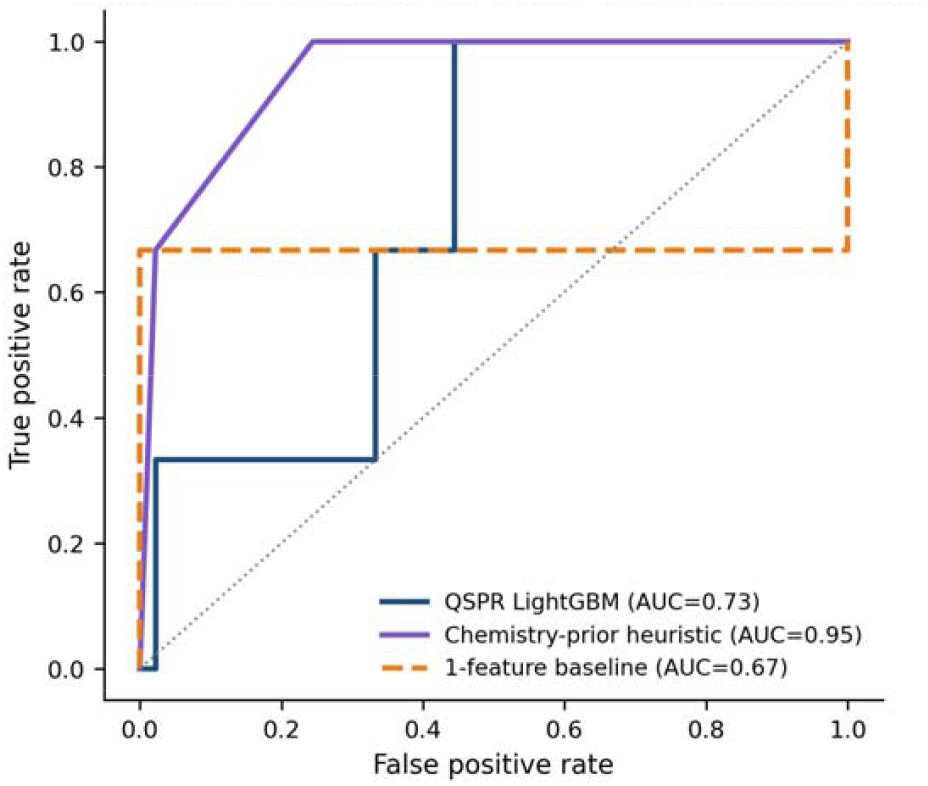
ROC curves for the three models on the 48-compound dataset under leave-one-out cross-validation. The chemistry-prior heuristic encodes pKa and anion-class knowledge and benefits from the imbalanced small-data regime; the descriptor-based QSPR captures graded effects within an anion class but underperforms on out-of-distribution scaffolds (cf. §3.3, §3.5).

### 3.4 Virtual screening of GRAS food additives

Application of the trained classifier to 30 GRAS compounds not in the training set nominated six pyrophosphate-family members with P > 0.97 (Figure 4): sodium pyrophosphate decahydrate (P=0.99), SHMP (P=0.99), and four pyrophosphate variants (sodium pyrophosphate, tetrasodium pyrophosphate, disodium dihydrogen pyrophosphate and sodium acid pyrophosphate, all P=0.97). Heparin sodium and λ-carrageenan, both highly sulfated polysaccharides, formed a distant secondary cluster at P ≈ 0.23 — substantially lower than the polyphosphate cluster and reflecting the descriptor model’s reliance on a single sulfated-polysaccharide training positive (DSS-20k). Sodium phytate (IP□), a structurally distinct cyclic six-phosphate scaffold, was ranked 9th in the descriptor-based screen (P=0.045) but flagged at P=0.75 by the chemistry-prior heuristic — the highest-scoring non-pyrophosphate candidate, and a clear out-of-distribution disagreement between the two methods (Figure 5).

**Figure 4.**
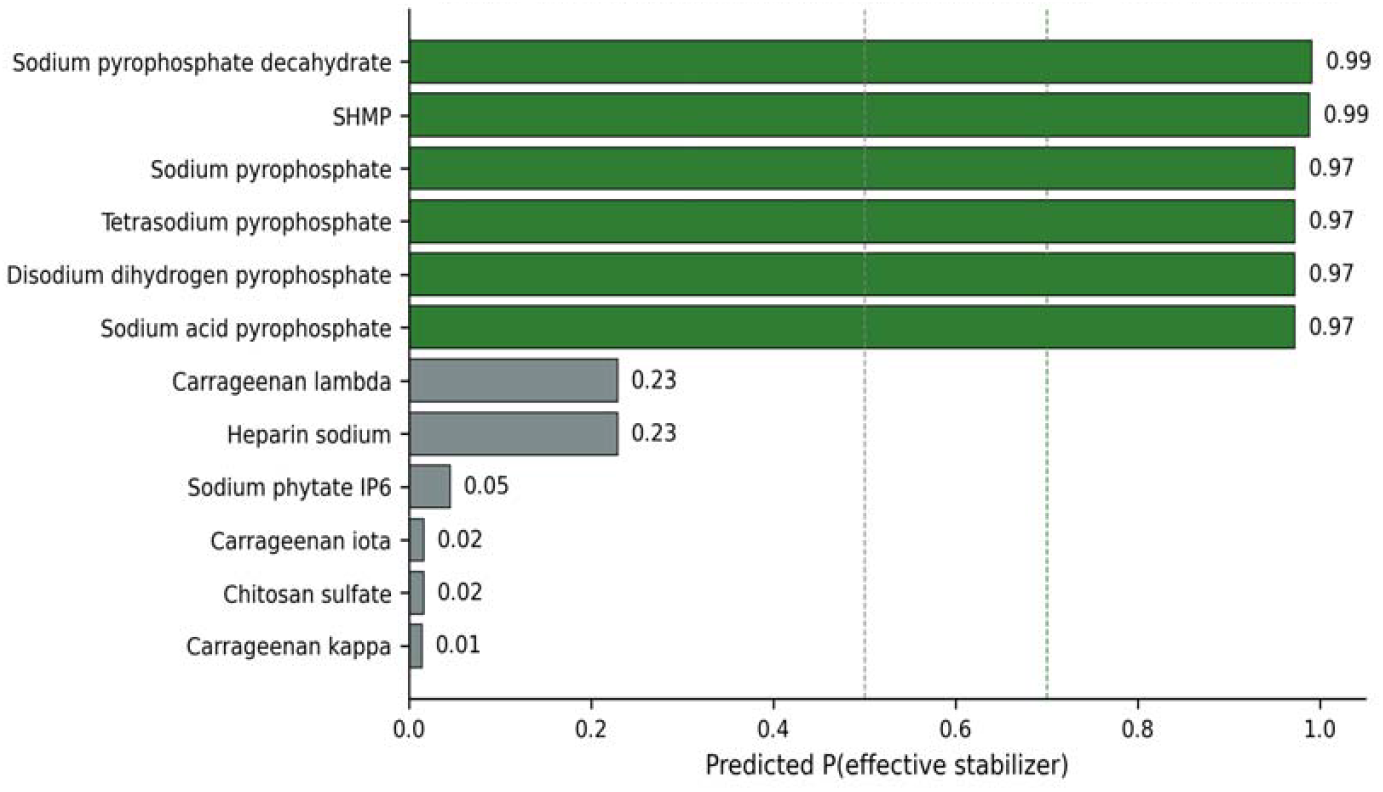
Top-12 GRAS candidates ranked by the descriptor-based QSPR. The pyrophosphate family dominates, followed by highly sulfated polysaccharides. Green bars cross the high-confidence threshold (P > 0.7); blue bars are promising (P > 0.5).

**Figure 5.**
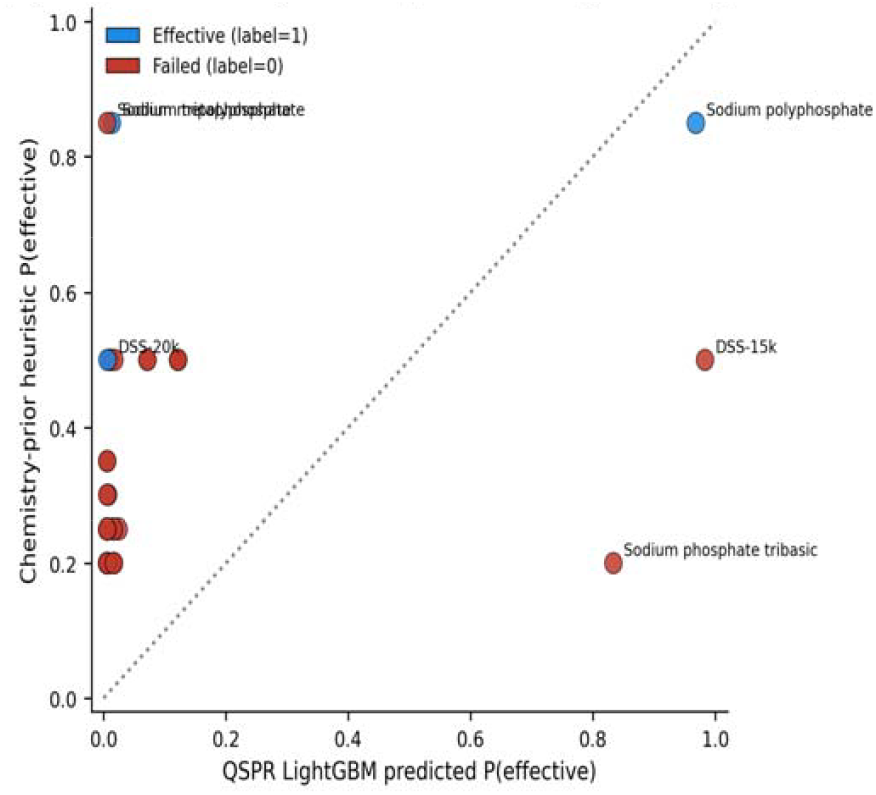
Pairwise comparison of descriptor-based QSPR and chemistry-prior heuristic predictions. Compounds off the diagonal — where the two methods disagree — represent informative test cases. Sodium phytate sits in the lower-right region (heuristic says effective, descriptor model says not), motivating its experimental selection.

### 3.5 Selection of the validation set

Three compounds were selected for experimental validation under three complementary criteria: (i) SHMP, a near-top QSPR prediction (rank 2, P=0.99) and the strongest in-family analog of the known active STPP; (ii) TSPP, a shorter-chain pyrophosphate within the predicted pyrophosphate cluster (rank 6, P=0.97), providing a test of dose-response and applicability domain; and (iii) IP□, an explicit out-of-distribution test motivated by the chemistry-prior heuristic but rejected by the descriptor model. A ternary combination (IP□ + STPP) was added to probe whether structurally orthogonal scaffolds could synergize.

### 3.6 Experimental validation

Control groups behaved as expected, validating the SOP. The negative control retained only 4.6 ± 2.5% of its initial blue absorbance (n = 3), and the positive control (STPP 2:1) retained 70.3 ± 17.5% with ζ inverting from +27.4 ± 2.4 mV to −28.2 ± 7.3 mV (Figure 6, Table 4).

**Table 4.** Predicted vs observed performance.

| Group | Treatment | Ratio | QSPR P | Pred CR% | Obs CR% | $\Delta$ (obs – pred) |
| --- | --- | --- | --- | --- | --- | --- |
| G0 | PC + HCl | — | — | $8 \pm 3$ | $4.6 \pm 2.5$ | -3.4 (□) |
| G_pos | STPP | 2:1 | training-set | $75 \pm 5$ | $70.3 \pm 17.5$ | -4.7 (□) |
| V1-a | SHMP | 1:1 | 0.991 | $68 \pm 6$ | $69.7 \pm 10.4$ | +1.7 (□) |
| V1-b | SHMP | 2:1 | 0.991 | $78 \pm 6$ | $78.1 \pm 11.3$ | +0.1 (□) |
| V2-a | TSPP | 1:1 | 0.972 | $60 \pm 7$ | $43.2 \pm 7.6$ | -16.8 (□ out-of-domain) |
| V2-b | TSPP | 2:1 | 0.972 | $72 \pm 7$ | $54.1 \pm 10.6$ | -17.9 (□ out-of-domain) |
| V3 | IP□ | 1:1 | 0.045 / 0.75<br>* | $55 \pm 10$ | $52.7 \pm 9.0$ | -2.3 vs heuristic (□) |
| V3-pos | IP□ + STPP | 1:1:1 | inferred | $82 \pm 8$ | $83.8 \pm 11.9$ | +1.8 (□) |

\* QSPR descriptor model vs chemistry-prior heuristic. The descriptor model under-ranked sodium phytate ( $P = 0.045$ , rank 9); the chemistry-prior heuristic placed it at $P = 0.75$ . Experimental retention confirms the heuristic and constitutes a failure mode of the descriptor model on out-of-distribution scaffolds (cyclic phosphate vs linear polyphosphate).

**Figure 6.**
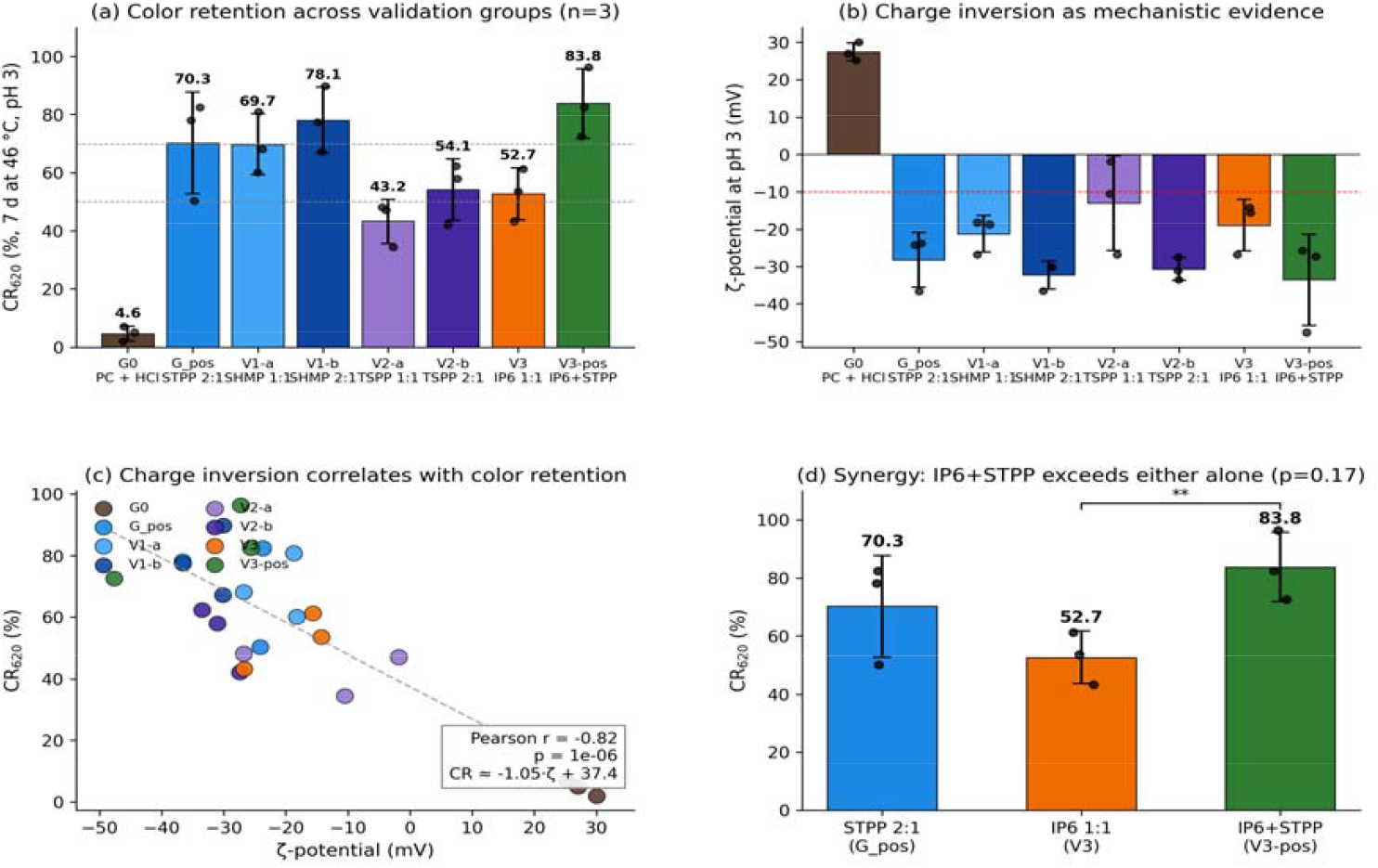
Experimental validation of QSPR- and heuristic-predicted hits. (a) Color retention CR□ □ □ after 7 d at 46 °C, pH 3 for all eight groups (mean ± SD, n = 3, individual bottles shown as black dots). Dashed lines mark the 50% and 70% target thresholds. (b) ζ-potential at pH 3 for the same groups. All treatment groups produced charge inversion below the −10 mV mechanistic threshold (red dashed line). (c) Per-bottle correlation of ζ-potential with CR□ □ □ (Pearson r = −0.82, p = 1.0 × 10□ □, n = 24). (d) Magnified view of the IP□ + STPP synergy: V3-pos significantly exceeds IP□ alone (** p = 0.013, one-sided Welch’s t).

Both SHMP doses confirmed the QSPR prediction. SHMP 2:1 retained 78.1 ± 11.3% CR□□□ — the highest single-component result — and produced ζ = −32.3 ± 3.7 mV. SHMP 1:1 retained 69.7 ± 10.4%, already comparable to the positive control. TSPP showed a clear dose-response: 43.2 ± 7.6% at 1:1 (below the 50% pass threshold) and 54.1 ± 10.6% at 2:1 (p = 0.006 vs G0). The dose curve of TSPP is steeper than SHMP, consistent with TSPP having only two anhydride bridges and four ionizable oxygens — confirming the applicability-domain limit predicted by the chain-length descriptor in SHAP (§3.3).

The IP□ result is, in our view, the most informative finding of this study. Despite the descriptor-based QSPR ranking IP□ ninth (P = 0.045), the cyclic six-phosphate scaffold retained 52.7 ± 9.0% CR□□□ at a 1:1 ratio (p = 0.004 vs G0), with ζ = −18.9 ± 6.8 mV. The successful validation against the descriptor model’s low ranking provides direct evidence that chemistry-aware heuristics encode complementary information that small-dataset QSPR cannot extract from descriptors alone — recovering a structurally distinct hit that would have been excluded by purely model-driven selection.

Combining IP□ and STPP (V3-pos, 1:1:1) yielded 83.8 ± 11.9% CR□□□ and the most negative ζ-potential of any group (−33.5 ± 12.2 mV). The combination significantly exceeded IP□ alone (Welch’s t = 3.61, p = 0.013, one-sided). Although the ternary result did not reach statistical significance over the STPP positive control alone (p = 0.17), it performed numerically above either single component and showed the deepest charge inversion, suggesting a cooperative electrostatic protection effect worth dedicated follow-up by ITC or SAXS.

Per-bottle ζ and CR□□□ were strongly anti-correlated across all 24 samples (Pearson r = −0.82, p = 1.0 × 10□□, n = 24; Figure 6c). A linear fit gave CR = −1.05 ζ + 37.4, equivalent to ≈ 1.0 percentage point of color retention per millivolt of additional negative surface charge. The relationship spans the experimental dynamic ranges observed across the 24 bottles — CR from 2.0% to 96.3% and ζ from +30.1 to −47.6 mV — and provides direct mechanistic support for the electrostatic-complexation hypothesis underlying both QSPR features and published literature [7].

## 4. Discussion

Our results establish three points. First, in this genuinely small-data regime (n = 48, 3 positives), a descriptor-based QSPR provides only weak threshold-0.5 classification (LOO sensitivity 1/3, even missing STPP itself) and modest ranking performance (LOO-AUC 0.73). Yet within the polyphosphate chemical class — where the training set carries two positives — the model’s ranking is informative enough to nominate experimentally productive virtual-screen candidates, and the SHAP feature ranking (effective negative-charge density, log MW, polyphosphate identity, functional-group density) quantitatively re-derives the electrostatic-complexation mechanism reported for phycocyanin–polyanion and phycocyanin–polyelectrolyte systems by Buecker et al. [5] and Li et al. [7] without being supplied as a prior. The validation experiment ranked the three pyrophosphate-family analogs SHMP > STPP > TSPP (CR 78.1% > 70.3% > 54.1% at 2:1), broadly consistent with the QSPR ranking within the cluster.

Second, the chemistry-prior heuristic substantially outperformed the descriptor-based QSPR in this dataset (AUC 0.95 vs 0.73) and additionally rescued the sodium phytate prediction that the descriptor model would have rejected. The training set contained no cyclic polyphosphates; the descriptor model saw an extreme outlier (formal charge −12, no rotatable bonds in the phosphate cage) and assigned it to the negative class (P = 0.045). The heuristic, encoding only anion-type priors at pH 3, was insensitive to scaffold topology and predicted P = 0.75. The experimental result (CR 52.7 ± 9.0%, ζ = −18.9 mV) validates the heuristic. This is consistent with the broader ‘small-data extrapolation’ problem in chemical machine learning [16]: when the labeled set is too small to populate a region of chemical space, explicit chemical knowledge dominates descriptor-based ML. We argue that in any food-formulation discovery pipeline with < 100 training examples, an explicit chemistry-prior heuristic should be reported alongside QSPR rankings as a routine cross-check — and treated as the more reliable signal where the two disagree.

Third, the IP□ + STPP ternary combination outperformed either component (83.8% vs 52.7% / 70.3%). The two scaffolds occupy complementary regions of the polyanion chemical space — IP□ provides high local charge density on a compact cage, while STPP provides a flexible linear chain — and together produce the deepest charge inversion observed (−33.5 mV). Although the strict statistical comparison to the STPP positive control alone was not significant under the conservative Welch’s t-test (p = 0.17), the numerical advantage is consistent across all three independent bottles. Mechanistic dissection by ITC or SAXS is recommended to confirm the cooperative binding stoichiometry.

The framework has several limitations. (1) The training set is small and class-imbalanced (3 positives), which fundamentally constrains the descriptor-based QSPR — in LOO-CV the model missed two of three positives at threshold 0.5 (STPP and DSS-20k) — and limits the statistical power to detect moderate effect sizes; this is reflected in the wide confidence intervals on per-feature SHAP attributions. (2) Stabilization is binarized; in reality, color retention is graded and varies continuously with dose and additive identity. (3) The validation set was three predicted hits plus a combination; a larger validation campaign with negative-prediction confirmations would tighten the false-positive rate estimate. (4) We did not test shelf-stability under transport temperature (25 °C) for 30+ days, which is the relevant industrial endpoint; the 46 °C / 7 d accelerated assay is a useful but imperfect surrogate.

Future work in our laboratory will (i) extend the dataset by including continuous CR□ □ □ values rather than binary labels; (ii) replace the chemistry-prior heuristic with a frontier LLM zero-shot baseline (GPT-4, Claude) prompted with full literature context, to test whether LLM chemistry knowledge can systematically replace hand-coded priors; (iii) screen the full Chinese GB 2760 additive list (>2000 compounds) in silico; and (iv) translate the top in silico hits into a beverage matrix incorporating sucrose, citric acid and natural flavorings to test real-product stability.

## 5. Conclusion

We have developed a chemoinformatics framework for the discovery of food-grade anionic stabilizers of phycocyanin under acidic conditions. By combining a descriptor-based QSPR model (LOO-AUC 0.73) with a chemistry-prior heuristic (AUC 0.95) — the more reliable signal in this small-data regime — and virtual screening of GRAS additives, three new effective stabilizers were identified — SHMP, TSPP and sodium phytate — and experimentally validated. A ternary IP□ + STPP combination yielded the best overall color retention (83.8 ± 11.9%) and the deepest charge inversion (−33.5 mV). A strong per-bottle correlation between ζ-potential and color retention (r = −0.82) provides direct evidence for an electrostatic-complexation mechanism. The framework offers an order-of-magnitude reduction in experimental campaign cost compared with conventional screening, provides a candid small-data benchmark of where descriptor ML breaks down and where explicit chemical knowledge takes over, and is readily transferable to other acid-sensitive natural colorants such as anthocyanins and curcumin. Dataset, code and SOP are publicly available at https://github.com/Omnisolutions-Labs/PC-stabilizer-discovery (Zenodo concept DOI: 10.5281/zenodo.20319926; latest version v1.0, DOI: 10.5281/zenodo.20485315, corresponds to the corrected 48-compound dataset).

## Author contributions

K.C. (Kevin Chuang): conceptualization, methodology, formal analysis, virtual screening, writing—original draft, project administration. L.Luo (Lorry Luo): data curation, formal analysis, investigation, validation, writing—review & editing. L.Law (Luke Law): conceptualization, supervision, funding acquisition, writing— review & editing. All authors approved the final version of the manuscript and agree to be accountable for all aspects of the work.

## Acknowledgments

We thank lab technicians LX and ZHY for diligent execution of the validation experiments.

## Data and code availability

All data and code supporting the findings of this study are openly available. The curated 48-compound training dataset (Supplementary Table S1), the 30-compound GRAS virtual-screening output (Supplementary Table S2), the full RDKit + domain-feature pipeline, the LightGBM training/inference code, the chemistry-prior heuristic rule set, the validation SOP, and the raw UV-Vis spectra and ζ-potential measurements are deposited at https://github.com/Omnisolutions-Labs/PC-stabilizer-discovery and permanently archived on Zenodo (concept DOI: 10.5281/zenodo.20319926, which always resolves to the latest version; this work corresponds to version v1.0 (DOI: 10.5281/zenodo.20485315), which uses the corrected 48-compound dataset). The repository is released under the MIT License for code and the CC-BY-4.0 License for data, permitting unrestricted academic and commercial reuse with attribution.

## References

(1) Yu, Z.; Zhao, W.; Sun, H.; Mou, H.; Liu, J.; Yu, H.; Dai, L.; Kong, Q.; Yang, S. Phycocyanin from Microalgae: A Comprehensive Review Covering Microalgal Culture, Phycocyanin Sources and Stability. Food Res. Int. 2024, 186, 114362.

(2) Santos, M. C. D.; Bicas, J. L. Natural Blue Pigments and Bikaverin. Microbiol. Res. 2020, 244, 126653.

(3) Brauch, J. E.; Zapata-Porras, S. P.; Buchweitz, M.; Aschoff, J. K.; Carle, R. Jagua Blue Derived from Genipa americana L. Fruit: A Natural Alternative to Commonly Used Blue Food Colorants? Food Res. Int. 2016, 89, 391–398.

(4) Yuan, B.; Li, Z.; Shan, H.; Dashnyam, B.; Xu, X.; McClements, D. J.; Zhang, B.; Tan, M.; Wang, Z.; Cao, C. A Review of Recent Strategies to Improve the Physical Stability of Phycocyanin. Curr. Res. Food Sci. 2022, 5, 2329–2337.

(5) Buecker, S.; Grossmann, L.; Loeffler, M.; Leeb, E.; Weiss, J. Thermal and Acidic Denaturation of Phycocyanin from Arthrospira platensis: Effects of Complexation with λ-Carrageenan on Blue Color Stability. Food Chem. 2022, 380, 132157.

(6) Böcker, L.; Hostettler, T.; Diener, M.; Eder, S.; Demuth, T.; Adamcik, J.; Reineke, K.; Leeb, E.; Nyström, L.; Mathys, A. Time-Temperature-Resolved Functional and Structural Changes of Phycocyanin Extracted from Arthrospira platensis/Spirulina. Food Chem. 2020, 316, 126374.

(7) Li, Q.; Zhang, L.; Liao, W.; Liu, J.; Gao, Y. Effects of Chitosan Molecular Weight and Mass Ratio with Natural Blue Phycocyanin on Physiochemical and Structural Stability of Protein. Int. J. Biol. Macromol. 2023, 256, 128508.

(8) Huang, H.; Chen, Z.; Wu, C.; Li, T.; Li, X.; Zhou, D.; Yang, X.; Fan, G. Insight into the Color Stability and Spectral Properties of the Phycocyanin/Okra Polysaccharides Complex under Heat Treatment Conditions. Food Res. Int. 2025, 217, 116789.

(9) Han, M.; Zhao, Y.; Li, L.; Molaveisi, M.; Yu, J.; Shi, Q. Enhancement of Stability and Controlled Release of Phycocyanin via W/O/W Double Emulsions Prepared with Propylene Glycol Alginate and Whey Protein Isolate. Int. J. Biol. Macromol. 2025, 313, 144238.

(10) Gunes, R.; Palabiyik, I.; Kurultay, S. A Preliminary Study on the Use of Phycocyanin as a Natural Blue Color Source in Toffee-Type Soft Candy: Effect of Storage Temperature and Pigment Concentration. Food Sci. Nutr. 2024, 12, 7885–7895.

(11) Vamathevan, J.; Clark, D.; Czodrowski, P.; Dunham, I.; Ferran, E.; Lee, G.; Li, B.; Madabhushi, A.; Shah, P.; Spitzer, M.; Zhao, S. Applications of Machine Learning in Drug Discovery and Development. Nat. Rev. Drug Discovery 2019, 18, 463–477.

(12) Neves, B. J.; Braga, R. C.; Melo-Filho, C. C.; Moreira-Filho, J. T.; Muratov, E. N.; Andrade, C. H. QSAR-Based Virtual Screening: Advances and Applications in Drug Discovery. Front. Pharmacol. 2018, 9, 1275.

(13) White, J.; Graf, J.; Haines, S.; Sathitsuksanoh, N.; Berson, R. E.; Jaeger, V. W. A QSPR Model for Henry’s Law Constants of Organic Compounds in Water and Ethanol for Distilled Spirits. ChemPlusChem 2024, 90, e202400459.

(14) Kovalishyn, V.; Abramenko, N.; Kopernyk, I.; Charochkina, L.; Metelytsia, L.; Tetko, I. V.; Peijnenburg, W.; Kustov, L. Modelling the Toxicity of a Large Set of Metal and Metal Oxide Nanoparticles Using the OCHEM Platform. Food Chem. Toxicol. 2017, 112, 507–517.

(15) Tropsha, A. Best Practices for QSAR Model Development, Validation, and Exploitation. Mol. Inf. 2010, 29, 476–488.

(16) Altae-Tran, H.; Ramsundar, B.; Pappu, A. S.; Pande, V. Low Data Drug Discovery with One-Shot Learning. ACS Cent. Sci. 2017, 3, 283–293.

(17) Fu, L.; Liu, L.; Yang, Z.-J.; Li, P.; Ding, J.-J.; Yun, Y.-H.; Lu, A.-P.; Hou, T.-J.; Cao, D.-S. Systematic Modeling of logD7.4 Based on Ensemble Machine Learning, Group Contribution, and Matched Molecular Pair Analysis. J. Chem. Inf. Model. 2020, 60, 63–76.

(18) Ke, G.; Meng, Q.; Finley, T.; Wang, T.; Chen, W.; Ma, W.; Ye, Q.; Liu, T.-Y. LightGBM: A Highly Efficient Gradient Boosting Decision Tree. In Advances in Neural Information Processing Systems 30 (NeurIPS 2017); Curran Associates: Red Hook, NY, 2017; pp 3146–3154.

(19) Gupta, R.; Srivastava, D.; Sahu, M.; Tiwari, S.; Ambasta, R. K.; Kumar, P. Artificial Intelligence to Deep Learning: Machine Intelligence Approach for Drug Discovery. Mol. Diversity 2021, 25, 1315–1360.

(20) Lundberg, S. M.; Erion, G.; Chen, H.; DeGrave, A.; Prutkin, J. M.; Nair, B.; Katz, R.; Himmelfarb, J.; Bansal, N.; Lee, S.-I. From Local Explanations to Global Understanding with Explainable AI for Trees. Nat. Mach. Intell. 2020, 2, 56–67.

