## Supplemental Data 1 for "Chemoinformatics-guided discovery of food-grade anionic stabilizers for phycocyanin under acidic conditions"

Manuscript version v3.0 · Compiled 2026-05-23

### **Contents**

• Table S1 — full 48-compound training dataset (page 2)

• Table S2 — 30-compound GRAS virtual-screening output (page 3)

• S3. Methods supplement — feature engineering, hyperparameters, SHAP, chemistry-prior heuristic, statistical and reproducibility notes (page 4)

• S4. Validation experiment Standard Operating Procedure (SOP) — pre-registered protocol for the 24-bottle validation campaign (page 8)

**Companion data files:** All raw and derived data files supporting this work (curated_compounds.csv, features_matrix.csv, training_predictions.csv, model_metrics.json, virtual_screening_results.csv, validation UV and ζ-potential per-bottle CSVs, and source code) are openly available at https://github.com/Omnisolutions-Labs/PC-stabilizer-discovery and permanently archived on Zenodo (concept DOI: 10.5281/zenodo.20319926).

### **Table S1 · Full 48-compound training dataset**

*The 48 unique compounds aggregated from three rounds of in-house screening (November 2025 – March 2026). Label ‘effective’ = preserved blue color across at least one PC ratio under the standardized condition (0.375% w/w PC, pH 3, 46 °C, 7 d); ‘failed’ = did not preserve color; ‘failed (ppt)’ = produced visible precipitate at the working concentration. Source-round date indicates the round in which the compound was last tested.*

| **#** | **Compound** | **Category** | **MW (g/mol)** | **Polymer** | **Anion class** | **FG density** | **Label** | **Source** |
| --- | --- | --- | --- | --- | --- | --- | --- | --- |
| 1 | Sodium chloride | salt | 58.44 | no | none | 0 | failed | 3.18 |
| 2 | Potassium chloride | salt | 74.55 | no | none | 0 | failed | 3.18 |
| 3 | Calcium chloride | salt | 110.98 | no | none | 0 | failed | 3.18 |
| 4 | Magnesium chloride | salt | 95.21 | no | none | 0 | failed | 3.18 |
| 5 | Zinc chloride | salt | 136.29 | no | none | 0 | failed | 3.18 |
| 6 | Ferric chloride | salt | 162.20 | no | none | 0 | failed (ppt) | 3.18 |
| 7 | Potassium sulfate | salt | 174.26 | no | sulfate | 2 | failed (ppt) | 3.18 |
| 8 | Calcium sulfate | salt | 136.14 | no | sulfate | 2 | failed (ppt) | 3.18 |
| 9 | Magnesium sulfate | salt | 120.37 | no | sulfate | 2 | failed (ppt) | 3.18 |
| 10 | Sodium phosphate tribasic | salt | 163.94 | no | phosphate | 3 | failed (ppt) | 3.18 |
| 11 | Sodium tripolyphosphate (STPP) | polyphosphate | 367.86 | no | polyphosphate | 5 | effective | 3.18 |
| 12 | Sodium polyphosphate | polyphosphate | 611.77 | yes | polyphosphate | 6 | effective | 11.24 |
| 13 | Sodium metaphosphate | polyphosphate | 101.96 | yes | metaphosphate | 1 | failed | 11.24 |
| 14 | Calcium alginate | polysaccharide | 200.00 | yes | carboxylate | 1 | failed (ppt) | 3.18 |
| 15 | Sodium alginate | polysaccharide | 198.00 | yes | carboxylate | 1 | failed | literature |
| 16 | Fucoidan | polysaccharide | 20000.00 | yes | sulfate | 1.2 | failed | 11.24 |
| 17 | Dextran sulfate Na 500k | polysaccharide | 500000.00 | yes | sulfate | 1.6 | failed | 11.24 |
| 18 | Dextran sulfate Na 100k | polysaccharide | 100000.00 | yes | sulfate | 1.6 | failed | 11.24 |
| 19 | Dextran sulfate Na 40k | polysaccharide | 40000.00 | yes | sulfate | 1.65 | failed | 11.24 |
| 20 | Dextran sulfate Na 20k | polysaccharide | 20000.00 | yes | sulfate | 1.7 | effective | 11.24 |
| 21 | Dextran sulfate Na 15k | polysaccharide | 15000.00 | yes | sulfate | 1.71 | failed | 11.24 |
| 22 | Dextran sulfate Na 5k | polysaccharide | 5000.00 | yes | sulfate | 1.78 | failed | 11.24 |
| 23 | Dextran sulfate Na 3k | polysaccharide | 3000.00 | yes | sulfate | 1.79 | failed | 11.24 |
| 24 | Lignosulfonate Na | polymer | 10000.00 | yes | sulfonate | 1 | failed | 11.24 |
| 25 | Sugammadex sodium | cyclodextrin | 2178.00 | no | carboxylate | 8 | failed | 1.30 |
| 26 | Glucose | sugar | 180.16 | no | hydroxyl | 5 | failed | 3.18 |
| 27 | Fructose | sugar | 180.16 | no | hydroxyl | 5 | failed | 3.18 |
| 28 | Sucrose | sugar | 342.30 | no | hydroxyl | 8 | failed | 3.18 |
| 29 | Trehalose | sugar | 342.30 | no | hydroxyl | 8 | failed | 3.18 |
| 30 | FOS (fructooligosaccharide) | sugar | 500.00 | yes | hydroxyl | 12 | failed | 3.18 |
| 31 | IMO (isomalto-oligosaccharide) | sugar | 500.00 | yes | hydroxyl | 12 | failed | 3.18 |
| 32 | GOS (galacto-oligosaccharide) | sugar | 500.00 | yes | hydroxyl | 12 | failed | 3.18 |
| 33 | Resistant starch | polysaccharide | 50000.00 | yes | hydroxyl | 3 | failed | 3.18 |
| 34 | Dextrin | polysaccharide | 20000.00 | yes | hydroxyl | 3 | failed | 3.18 |
| 35 | Cellulose | polysaccharide | 100000.00 | yes | hydroxyl | 3 | failed | 3.18 |
| 36 | Inulin | polysaccharide | 5500.00 | yes | hydroxyl | 3 | failed | 3.18 |
| 37 | Pectin | polysaccharide | 150000.00 | yes | carboxylate | 0.7 | failed | 3.18 |
| 38 | Arabic gum | polysaccharide | 350000.00 | yes | carboxylate | 0.2 | failed | 3.18 |
| 39 | Xylitol | sugar_alcohol | 152.15 | no | hydroxyl | 5 | failed | 3.18 |
| 40 | Sorbitol | sugar_alcohol | 182.17 | no | hydroxyl | 6 | failed | 3.18 |
| 41 | Acesulfame potassium | sweetener | 201.24 | no | sulfonate | 1 | failed (ppt) | 3.18 |
| 42 | Citric acid | organic_acid | 192.12 | no | carboxylate | 3 | failed | 3.18 |
| 43 | Malic acid | organic_acid | 134.09 | no | carboxylate | 2 | failed | 3.18 |
| 44 | Lactic acid | organic_acid | 90.08 | no | carboxylate | 1 | failed | 3.18 |
| 45 | Tartaric acid | organic_acid | 150.09 | no | carboxylate | 2 | failed | 3.18 |
| 46 | Ascorbic acid | organic_acid | 176.12 | no | carboxylate | 2 | failed | 3.18 |
| 47 | Glycerol | other | 92.09 | no | hydroxyl | 3 | failed | 3.18 |
| 48 | Gelatin | protein | 50000.00 | yes | amide | 0 | failed (ppt) | 3.18 |

### **Table S2 · Virtual-screening output (30 GRAS food additives)**

*Compounds not present in the training set, scored by the final LightGBM QSPR classifier (retrained on all 48 training compounds). Ranked by descriptor-based P(effective). The chemistry-prior heuristic separately assigned P=0.75 to sodium phytate (rank 9 in this table) and P=0.85 to the six pyrophosphate-family entries — see manuscript §3.4 and Figure 5.*

| **Rank** | **Compound** | **Anion class** | **Polymer** | **MW (g/mol)** | **FG density** | **P(effective)** | **Recommendation** |
| --- | --- | --- | --- | --- | --- | --- | --- |
| 1 | Sodium pyrophosphate decahydrate | polyphosphate | no | 446.06 | 4.0 | 0.991 | ★★ Strong |
| 2 | Sodium hexamethaphosphate (SHMP) | polyphosphate | yes | 611.77 | 6.0 | 0.991 | ★★ Strong |
| 3 | Sodium pyrophosphate | polyphosphate | no | 265.9 | 4.0 | 0.972 | ★★ Strong |
| 4 | Disodium dihydrogen pyrophosphate | polyphosphate | no | 221.9 | 2.0 | 0.972 | ★★ Strong |
| 5 | Sodium acid pyrophosphate | polyphosphate | no | 221.9 | 2.0 | 0.972 | ★★ Strong |
| 6 | Tetrasodium pyrophosphate | polyphosphate | no | 265.9 | 4.0 | 0.972 | ★★ Strong |
| 7 | Heparin sodium | sulfate | yes | 15000.0 | 2.7 | 0.229 | — |
| 8 | Carrageenan lambda | sulfate | yes | 400000.0 | 2.4 | 0.229 | — |
| 9 | Sodium phytate (IP6) | phosphate | no | 660.0 | 6.0 | 0.045 | — |
| 10 | Chitosan sulfate | sulfate | yes | 20000.0 | 1.5 | 0.016 | — |
| 11 | Carrageenan iota | sulfate | yes | 400000.0 | 1.6 | 0.016 | — |
| 12 | CMC (carboxymethyl cellulose) | carboxylate | yes | 250000.0 | 0.7 | 0.014 | — |
| 13 | Gellan gum | carboxylate | yes | 500000.0 | 0.5 | 0.014 | — |
| 14 | Sodium dextran phosphate | phosphate | yes | 40000.0 | 1.2 | 0.014 | — |
| 15 | Carrageenan furcellaran | sulfate | yes | 100000.0 | 0.8 | 0.014 | — |
| 16 | Locust bean gum | hydroxyl | yes | 300000.0 | 3.0 | 0.014 | — |
| 17 | Konjac glucomannan | hydroxyl | yes | 1000000.0 | 3.0 | 0.014 | — |
| 18 | Whey protein isolate | carboxylate | yes | 18000.0 | 0.3 | 0.014 | — |
| 19 | Sodium caseinate | carboxylate | yes | 24000.0 | 0.3 | 0.014 | — |
| 20 | Dipotassium phosphate | phosphate | no | 174.18 | 2.0 | 0.014 | — |
| 21 | Carrageenan kappa | sulfate | yes | 400000.0 | 0.8 | 0.014 | — |
| 22 | Pectin (low methoxyl) | carboxylate | yes | 150000.0 | 0.7 | 0.014 | — |
| 23 | Xanthan gum | carboxylate | yes | 2000000.0 | 0.5 | 0.014 | — |
| 24 | Alginic acid (low Mw) | carboxylate | yes | 30000.0 | 1.0 | 0.014 | — |
| 25 | Hyaluronic acid | carboxylate | yes | 1000000.0 | 0.5 | 0.014 | — |
| 26 | Chondroitin sulfate | sulfate | yes | 30000.0 | 1.0 | 0.014 | — |
| 27 | Pectin (high methoxyl) | carboxylate | yes | 150000.0 | 0.3 | 0.014 | — |
| 28 | Sodium ascorbate | carboxylate | no | 198.1 | 1.0 | 0.006 | — |
| 29 | Trisodium citrate | carboxylate | no | 258.1 | 3.0 | 0.006 | — |
| 30 | Polylysine epsilon | amide | yes | 4000.0 | 0.0 | 0.006 | — |

### **Methods supplement**

This document expands on the Methods section of the manuscript with implementation details, parameter rationales, and reproducibility notes.

#### **A. Feature engineering**

The 21-dimensional feature vector combines 10 RDKit descriptors and 11 hand-curated domain-expert features.

##### ***A.1 RDKit descriptors (10)***

| **Feature** | **RDKit call** | **Notes** |
| --- | --- | --- |
| MolWt | `Descriptors.MolWt(mol)` | g/mol |
| LogP | `Crippen.MolLogP(mol)` | dimensionless |
| TPSA | `Descriptors.TPSA(mol)` | Å² |
| HBD | `Lipinski.NumHDonors(mol)` | count |
| HBA | `Lipinski.NumHAcceptors(mol)` | count |
| FormalCharge | sum of `GetFormalCharge()` over atoms | integer |
| NumRings | `Lipinski.RingCount(mol)` | count |
| NumRotBonds | `Lipinski.NumRotatableBonds(mol)` | count |
| FracSP3 | `rdMolDescriptors.CalcFractionCSP3(mol)` | 0–1 |
| NumHeavyAtoms | `mol.GetNumHeavyAtoms()` | count |

Polymer compounds without a canonical SMILES are assigned default RDKit values:

MolWt=mw, LogP=-2.0, TPSA=200, HBD=10, HBA=15,
FormalCharge=-int(eff_neg_charge_per_unit*5),
NumRings=1, NumRotBonds=20, FracSP3=0.9, NumHeavyAtoms=50

##### ***A.2 Domain-expert features (11)***

| **Feature** | **Definition** |
| --- | --- |
| is_polymer | 0/1 |
| log10_mw | log₁₀ of molecular weight |
| is_phosphate | 1 if anion class ∈ {phosphate, polyphosphate, metaphosphate} |
| is_polyphosphate | 1 if anion class = polyphosphate |
| is_sulfate / is_sulfonate / is_carboxylate | one-hot encoding |
| is_anionic | 1 if any anionic group present |
| fg_density | functional groups per repeat unit |
| eff_neg_charge_per_unit | fg_density × pKa-weighted ionization fraction at pH 3 |
| is_metal_polyvalent | 1 if compound contains Ca²⁺, Mg²⁺, Zn²⁺, Fe³⁺, etc. |

The effective negative charge weighting at pH 3 uses:

• Polyphosphate, sulfate, sulfonate: 1.0 (fully ionized)

• Metaphosphate: 0.7

• Carboxylate: 0.4 (partial ionization; pKa ≈ 3–4)

• Hydroxyl, amide, none: 0.0

#### **B. Model hyperparameters and training**

LightGBM classifier:

```
LGBMClassifier(
    n_estimators=200,
    learning_rate=0.04,
    num_leaves=15,
    min_child_samples=2,
    max_depth=4,
    reg_lambda=1.0,
    class_weight={0: 1.0, 1: pos_weight},  # pos_weight = n_neg / n_pos
    random_state=20260519
)
```

`pos_weight` is computed dynamically as `(n_total − n_positive) / n_positive` = 15 for the corrected 48-compound dataset (3 positives, 45 negatives). Leave-one-out cross-validation is used due to the small dataset size; 48 LightGBM models are trained, each on 47 examples.

#### **C. SHAP attribution**

`shap.TreeExplainer` is applied to the final model (trained on all 48 examples) to obtain Shapley values for each feature on each compound. The summary plot reflects the absolute Shapley contributions across the full dataset.

#### **D. Chemistry-prior heuristic**

The chemistry-prior heuristic is a deterministic rule set assigning P(effective) directly from compound name and anion class (see `code/qspr_pipeline.py`, function `chem_prior` / `llm_like_zero_shot`). It encodes published pKa values and reported activity of major anion classes at pH 3.

Probabilities are calibrated against the 48-compound training set; refinement against an external benchmark dataset is left to future work.

#### **E. Virtual screening library**

Thirty GRAS food additives were curated by combining EFSA E-number lists with the Chinese GB 2760 additive catalog, restricted to anionic polymers and phosphates. Polymers without canonical SMILES use the polymer default descriptor set described in §A.1.

#### **F. Statistical analysis details**

• One-sided Welch's t-test (`scipy.stats.ttest_ind(..., equal_var=False, alternative='greater')`) for treatment vs G0 contrasts.

• Pearson r and two-tailed p-value via `scipy.stats.pearsonr` for ζ vs CR₆₂₀.

• Holm-Bonferroni correction across the seven vs-G0 contrasts at family-wise α = 0.05.

#### **G. Reproducibility**

Random seed: `SEED = 20260519`. All NumPy random generators, LightGBM `random_state`, and Monte-Carlo sampling routines use this seed. Re-running `python code/qspr_pipeline.py` and `python code/analyze_real.py` reproduces all reported AUC, accuracy, virtual-screening ranks, and Figure 6 panels at bit-level (verified on Python 3.13, numpy 1.26, pandas 2.0, scipy 1.15.3, scikit-learn 1.5, lightgbm 4.0, shap 0.45).

#### **H. Limitations**

1. Class imbalance (3 positives / 45 negatives) fundamentally limits LOO-CV power; SHAP attributions are interpreted qualitatively.

2. Polymer compounds with default descriptors may underestimate within-class differences.

3. Sodium phytate (IP₆) was a known descriptor-model false negative; the model's applicability domain is therefore limited to linear polyphosphates and acyclic polyanions.

4. The 7-day accelerated assay at 46 °C is a surrogate for the industrially relevant 30-day shelf life at 25 °C; real-product testing is outside the present scope.

### **Validation Experiment Standard Operating Procedure (SOP)**

*Companion to: *Chemoinformatics-guided discovery of food-grade anionic stabilizers for phycocyanin under acidic conditions* (Chuang et al., 2026).*

This SOP corresponds to the 24-bottle wet-lab validation experiment reported in Section 3.6 of the manuscript. All quantitative data from this protocol are released in `data/validation/`.

#### **1. Scope and acceptance criteria**

Validate the QSPR- and chemistry-prior-predicted top candidates (SHMP, TSPP, sodium phytate) at pH 3 and 46 °C over 7 days.

Acceptance criteria (any one yields a primary result):

• SHMP (V1-a or V1-b) CR₆₂₀ ≥ 60%

• TSPP (V2-a or V2-b) CR₆₂₀ ≥ 50%

• Sodium phytate (V3) CR₆₂₀ ≥ 50%

• ζ-potential of any treatment group at pH 3 below −10 mV

#### **2. Experimental matrix (24 bottles, n = 3)**

| **Group** | **Treatment** | **Ratio (stab:PC)** | **n** | **Bottle IDs** |
| --- | --- | --- | --- | --- |
| G0 | PC + HCl (negative) | — | 3 | B01–B03 |
| G_pos | STPP (positive control) | 2:1 | 3 | B04–B06 |
| V1-a | SHMP (top-1 QSPR) | 1:1 | 3 | B07–B09 |
| V1-b | SHMP | 2:1 | 3 | B10–B12 |
| V2-a | TSPP | 1:1 | 3 | B13–B15 |
| V2-b | TSPP | 2:1 | 3 | B16–B18 |
| V3 | Sodium phytate (IP₆) | 1:1 | 3 | B19–B21 |
| V3-pos | IP₆ + STPP (ternary) | 1:1:1 | 3 | B22–B24 |

#### **3. Critical control points (CCP)**

| **CCP** | **Parameter** | **Locked value** | **Failure mode if violated** |
| --- | --- | --- | --- |
| CCP-1 | PC final concentration | 0.375 % w/w | Baseline drift; CR not comparable |
| CCP-2 | Target pH | 3.00 ± 0.05 | ζ-titration inflection shifts ± 5 mV |
| CCP-3 | Incubation temp × time | 46.0 ± 0.5 °C × 168 h | Kinetics drift |
| CCP-4 | Sample / headspace | 10.0 mL / 14 mL | Oxidation accelerates fading |
| CCP-5 | PC stock thaw/refreeze | Single use, same day | Partial denaturation |
| CCP-6 | HCl titration rate | 1.00 M, 1 µL/s, 200 rpm stir | Local pH < 2 precipitation |

#### **4. Materials**

Food-grade phycocyanin E18+ (Yunnan Lü-A Bio, lot PC-E18-260509-B), STPP (Aladdin, lot STPP-260428-03), SHMP (Macklin, lot SHMP-260502-11), TSPP (Macklin, lot TSPP-260430-07), sodium phytate (Sigma, lot IP6-260421-02), 1.00 M HCl (titrated), 24 mL amber headspace vials, 0.22 µm PVDF filters, NIST-traceable pH calibration buffers.

#### **5. Workflow (10-day timeline)**

• **D-2** Materials ready; vials cleaned, dried, and labelled (B01–B24).

• **D-1** Stocks prepared: 0.50% w/w PC, 75 mg/mL stabilizer master solutions, 1 mM NaCl pH 3 ζ-dilution buffer; HOBO logger placed in incubator 24 h preheat.

• **D0** Sample preparation, t = 0 UV-Vis scan, N₂ purge, seal, incubate.

• **D1–D6** Twice-daily temperature check (09:00 / 17:00); HOBO logger continuous.

• **D7** End-point: ambient equilibration 30 min → UV-Vis full scan → D65 light-box photo → ζ-potential (1:10 dilution into 1 mM NaCl pH 3; 3 within-instrument repeats; latex standard every 5 samples).

• **D8** Data entry, blank correction, CR₆₂₀ calculation.

• **D9** Statistical analysis (Welch's t-tests vs G0; Pearson ζ-CR correlation).

#### **6. Sample preparation (per bottle, 10.0 mL final volume)**

| **Group** | **PC master (0.50%)** | **Stabilizer master (10×)** | **1 M HCl** | **ddH₂O to** |
| --- | --- | --- | --- | --- |
| G0 | 7.50 mL | — | ≈ 0.10 mL | 10.00 mL |
| G_pos | 7.50 mL | 1.00 mL STPP (75 mg/mL) | ≈ 0.13 mL | 10.00 mL |
| V1-a | 7.50 mL | 0.50 mL SHMP | ≈ 0.13 mL | 10.00 mL |
| V1-b | 7.50 mL | 1.00 mL SHMP | ≈ 0.13 mL | 10.00 mL |
| V2-a | 7.50 mL | 0.50 mL TSPP | ≈ 0.13 mL | 10.00 mL |
| V2-b | 7.50 mL | 1.00 mL TSPP | ≈ 0.13 mL | 10.00 mL |
| V3 | 7.50 mL | 1.00 mL IP₆ (37.5 mg/mL) | ≈ 0.15 mL | 10.00 mL |
| V3-pos | 7.50 mL | 0.5 mL IP₆ + 0.5 mL STPP | ≈ 0.15 mL | 10.00 mL |

#### **7. Measurement protocols**

**UV-Vis** — 1 cm quartz cell; spectrum 350–750 nm, step 1 nm, medium speed; blank = matching pH and stabilizer concentration without PC. CR₆₂₀ = [A₆₂₀(7d) − A_blank] / [A₆₂₀(0) − A_blank] × 100.

**ζ-Potential** — Malvern Zetasizer Nano ZS with DTS1070 folded-capillary cells; samples diluted 1:10 (or 1:20 if PDI > 0.5) in 1 mM NaCl, pH 3.00; 25 °C; 60 s equilibration; three runs per bottle. Latex standard target −55 ± 5 mV verified every five samples.

**Photo documentation** — D65 light box, ISO 100, f/8, 1/125 s, 50 mm; gray-card white balance; Pantone 285/286/2935 blue color cards as reference.

#### **8. Statistical analysis**

• One-sided Welch's t-test vs G0 for each treatment (H₁: treatment > G0)

• Pearson r between per-bottle ζ-potential and CR₆₂₀

• Holm-Bonferroni adjustment across the seven vs-G0 contrasts at family-wise α = 0.05

• Synergy test: one-sided Welch's t-test of V3-pos vs V3 alone

#### **9. Pre-registration**

This SOP was finalized and recorded internally on 2026-05-11 prior to the start of bottle preparation (2026-05-12), serving as a preregistration document for the validation campaign.
